# Meditope-Enabled Chimeric Antigen Receptors Facilitate Plug-and-Play Control of T Cells

**DOI:** 10.64898/2026.04.02.715976

**Authors:** Cheng-Fu Kuo, Zhen Tong, Yi-Chiu Kuo, Miso Park, Jeremy King, Kevin Ly, Bea Parcutela, Lawrence A. Stern, Zhiqiang Wang, Brenda Aguilar, Renate Starr, Wen-Chung Chang, Julie R. Ostberg, Daniel Rossi, Mary C. Clark, Darya Alizadeh, Stephen J. Forman, John C. Williams, Christine E. Brown

**Affiliations:** Department of Hematology and Hematopoietic Cell Transplantation, City of Hope, Duarte, CA; Irell and Manella Graduate School of Biological Sciences, City of Hope, Duarte, CA; Department of Cancer Biology and Molecular Medicine, Beckman Research Institute at City of Hope, Duarte, CA; Department of Clinical and Translational Project Development, City of Hope, Duarte, CA; Department of Chemical, Biological, and Materials Engineering, University of South Florida, Tampa, FL

## Abstract

Chimeric antigen receptor (CAR) T cells have transformed cancer treatment, yet challenges for achieving broader clinical success remain, including overcoming tumor antigen heterogeneity and limited T cell fitness. To address these challenges and enhance CAR T cell functionality, we leveraged meditope technology, a lock-and-key platform where Fab regions of antibodies are modified to bind a small cyclic peptide termed meditope (meP). We developed a panel of meditope-enabled Fab-based CARs (meCARs), which show selective binding to the meP and comparable activity to traditional single-chain variable fragment (scFv)-based CARs. Focusing on HER2-targeted meCARs for evaluating platform utility, we exploited the modularity of the meditope platform to detect meCAR T cells using meP-fused fluorescent agents, promote meCAR T cell expansion via meP-fused IL-15 cytokine, and broaden tumor antigen targeting through meP-fused antibodies to address tumor heterogeneity. These findings establish the meditope technology as a versatile strategy to augment CAR T cell functionality and overcome key limitations of current CAR-based therapies.

---

The performance of current chimeric antigen receptor (CAR) T cell therapies is constrained by fixed features encoded in the CAR design, limiting opportunities for fine-tuned regulation and expanded functionality. These hard-wired CARs have achieved remarkable clinical success in relapsed/refractory B-cell malignancies^1-5^, clinical settings where tumor antigen expression is relatively homogenous. However, these fixed CAR designs have had limited success against solid tumors, due to many challenges, including tumor heterogeneity, limited persistence, and the suppressive tumor microenvironment (TME). Further, unregulated CAR T cell expansion and cytokine production can result in life-threatening toxicities, including cytokine release syndrome and immune effector cell-associated neurotoxicity syndrome^6, 7^. These limitations underscore the need for more advanced CAR designs that enable tunable control and expanded functionality.

We incorporated meditope technology as part of a programmable CAR T cell system. Meditope technology effectively transforms a unique cyclic peptide (meditope peptide [meP]) and an engineered binding site within the fragment antigen-binding (Fab) region of an antibody (meditope-binding site) into an orthogonal but inert ligand/receptor pair that can be exploited to add functionality to any meditope-enabled Fab/monoclonal antibody (mAb). meP is a 12-amino acid cyclic peptide (sequence: CQFDLSTRRLKC) that was originally identified by its binding to the therapeutic antibody cetuximab through a noncovalent interaction without interfering with antigen recognition^8, 9^. Based on the crystal structure of the meP-cetuximab complex, distinct residues were identified and grafted onto human mAbs, enabling meP binding without affecting antigen binding affinity^8, 10^. Other compounds including fluorophores and biologics have been genetically or chemically coupled to meP, adding novel functionality to meditope-enabled (me) mAbs^9^.

We hypothesized that Fab-derived CARs could be meditope-enabled (meCAR) to allow binding of meP-fused compounds, offering a modular approach to enable CAR T cell programmability. As proof-of-concept, we successfully generated meCAR constructs by replacing the conventional single-chain variable fragment (scFv) domain with a meditope-enabled Fab. Furthermore, we utilized meP-fused compounds to visualize meCAR T cells by flow cytometry and immunofluorescence, enhance expansion of meCAR T cells and establish a dual-targeting meCAR T cell platform without additional genetic engineering. Our study demonstrates the potential of combining CAR T cell therapy with meditope technology to overcome limitations of current hard-wired CARs and advance the field of engineered cell therapies.

## Results

### Development of meCAR T cell platform

Unlike traditional CAR constructs that target via an scFv, engineering the meP binding site required construction of Fab-based CARs (**Fig. 1a**). We optimized Fab-CAR design using a trastuzumab-based HER2-targeted CAR, based on prior studies establishing the amino acid substitutions conferring meP binding^8^ (**Extended Data Fig. 1a**) and our experience with HER2-CAR T cells^11^. We optimized HER2-meCAR design, generating three constructs that varied in the meFab structural framework (**Fig. 1a**): 1) meCAR-LH (Fab light chain CAR, followed by T2A and Fab heavy chain), 2) meCAR-HL (Fab heavy chain CAR, followed by T2A and Fab light chain), and 3) meCAR-SC (single-chain (SC) Fab^12^ CAR). We confirmed meCAR expression in transduced healthy human donor T cells (**Fig. 1b**). Binding of meP-fused Alexa647 (meP-647) to meCAR T cells confirmed proper Fab folding and retention of the meP-binding site (**Fig. 1b**). The meCAR-SC construct displayed the highest CAR expression and meP-binding (**Fig. 1b**), as well as the strongest HER2 antigen binding (EC50 < 1.0 nM) (**Extended Data Fig. 1b**).

**Fig. 1:**
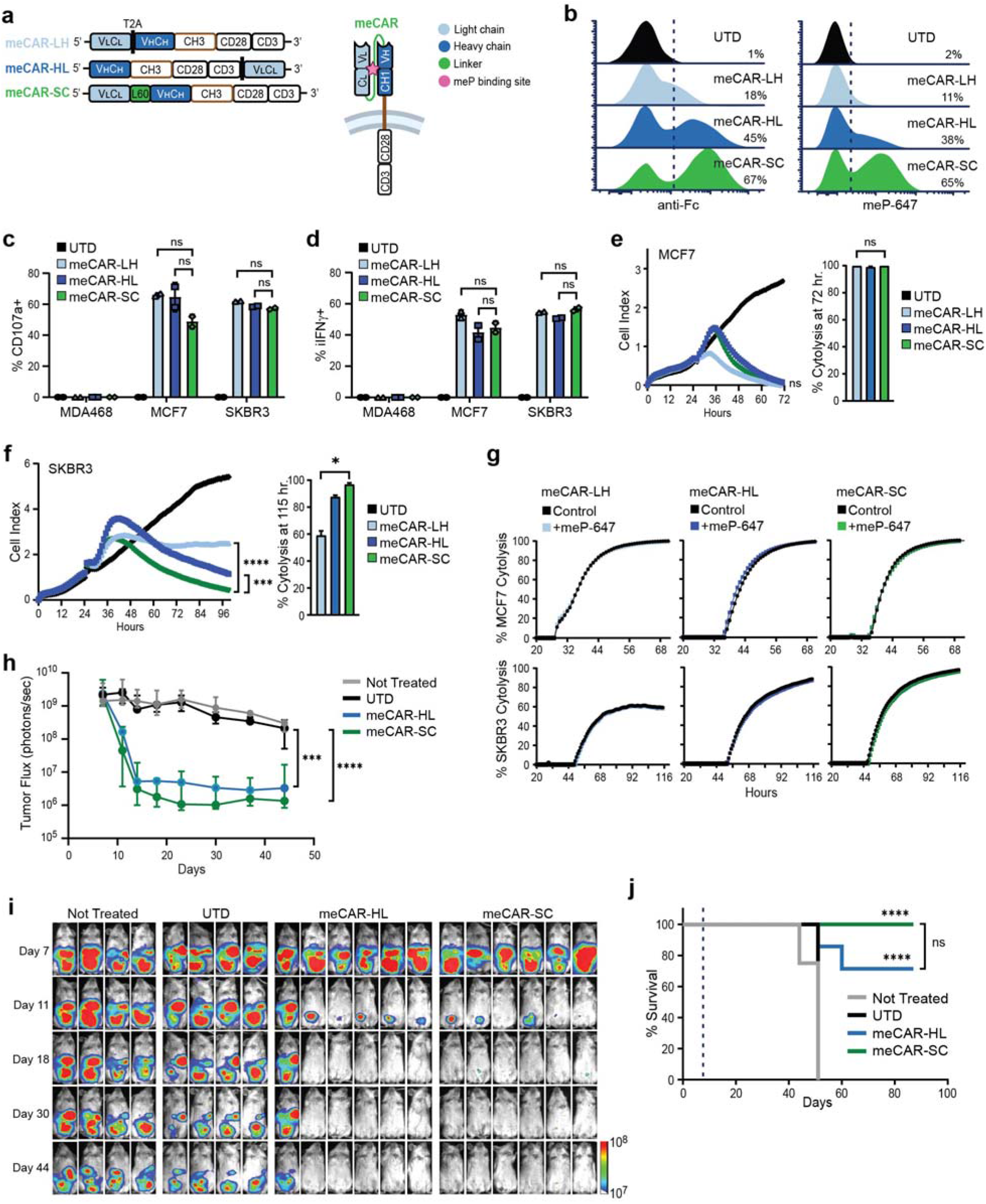
Engineering and characterization of meCAR T cells targeting HER2. **a**, Schematic of meCAR constructs, with light chain (V_L_C_L_) and heavy chain (V_H_C_H_) sequences depicted in relation to the IgG4-derived CH3 hinge, CD28 transmembrane/costimulatory, and CD3ζ signaling sequences, as well as the T2A ribosomal skip or L60 linker sequences as indicated. The meditope peptide (meP) binding domain is depicted in the resulting CAR protein schematic. **b**, meCAR expression by flow cytometric staining with anti-Fc (left) and meP binding using Alexa 647-fused meP (meP-647, right). Untransduced (UTD) cells were used as the negative control. **c**,**d**, Percentage of cells expressing CD107a (**c**) and intracellular IFNγ (**d**) for the indicated HER2-meCAR T cell variants (gated on viable CAR+ cells) after co-culture with HER2-negative MDA468 tumor cells, HER2-positive MCF7 cells or SKBR3 tumor cells using a one-way ANOVA test. **e**,**f**, Kinetics of tumor cell killing performed by xCELLigence assay upon co-culture of the indicated meCAR T cells with MCF7 (**e**) or SKBR3 (**f**) tumor cells. Mean cell index measurements reflecting tumor cell viability of triplicate wells at each time point are shown. Bar charts depict the percent cytolysis as compared to co-culture with UTD cells at last measured time point using a one-way ANOVA test. **g**, xCELLigence killing assays in the presence or absence of 100nM meP-647. Percent MCF7 (top) or SKBR3 (bottom) cytolysis upon co-culture with the indicated meCAR T cells are depicted at each time point. **h-j**, ffLuc+ MCF7 cells were intraperitoneally (i.p.) injected into NSG mice. On day 8 post-implantation, mice were left untreated or were treated with an i.p. injection of either UTD, meCAR-HL or meCAR-SC T cells followed by biophotonic imaging for tumor size over time (**h**,**i**). **h**, Mean ± S.D. tumor flux value is depicted for each group using a two-way ANOVA test. **j**, Kaplan-Meier survival analysis of each group using a Mantel-Cox test. Dashed line indicates day of T cell treatment. ns, non-significant *, P value <0.05; ***, P value <0.001; ****, P value <0.0001

We examined effector function of HER2-meCAR T cells by measuring CD107a degranulation and IFNγ secretion upon co-culture with MCF7 and SKBR3 breast cancer cell lines, which exhibit moderate and high HER2 expression (**Extended Data Fig. 2a**), respectively. We observed comparable activity for all meCAR T cell variants (**Fig. 1c, d**). In xCELLigence-based killing assays, HER2-meCAR T cells killed both MCF7 (**Fig. 1e**) and SKBR3 (**Fig. 1f**), with the meCAR-SC construct exhibiting superior killing of SKBR3 (**Fig. 1f**). Addition of meP-647 did not impact the ability of meCAR T cells to kill either cell line (**Fig. 1g**), confirming that binding of meP to meFab does not interfere with meCAR antigen recognition^8^. We compared in vivo antitumor activity of HER2-meCAR T cells, testing only the meCAR-HL and meCAR-SC constructs, which showed the highest meCAR expression (**Fig. 1b**) and *in vitro* functionality (**Fig. 1f**). Using an MCF7 tumor xenograft model in NSG mice, we observed potent antitumor activity in tumor-bearing mice treated with meCAR-HL or meCAR-SC compared to untreated or untransduced (UTD) T cell treated controls (**Fig. 1h, i**). The meCAR-SC T cells eliminated tumor in all mice (N=6) and prolonged survival for >80 days, modestly outperforming meCAR-HL (Fig. 1h-j). Moreover, meCAR-SC had comparable efficacy to our clinical scFv-based HER2-CAR^11^ in a HER2+ ovarian cancer xenograft model (**Extended Data Fig. 2 b-d**).

To evaluate applicability of the meCAR platform to other targets, we incorporated the SC-Fab framework and meditope-enabled two other CARs: 1) a humanized CD19-CAR recently developed by our group^13^; and 2) a HER3/EGFR dual-targeting CAR based on duligotuzumab (MEHD7945A)^14^. We confirmed meCAR-SC expression in T cells, binding of meP, and antigen-dependent CAR effector function (**Extended Data Fig. 3**). These results suggest that meCAR is a generalizable platform.

Overall, these findings demonstrated feasibility of generating Fab-based meCARs that bind meP without compromising CAR-mediated antigen binding and identified the meCAR-SC construct as the preferred meCAR design. Therefore, we moved forward with evaluating the utility of the meCAR platform, focusing on HER2-meCAR-SC as proof-of-concept.

### *In situ* detection of meCAR T cells

We evaluated the utility of the meditope technology for CAR T cell detection using fluorescent-fused mePs as tools for visualizing meCAR T cells by fluorescence microscopy. We stained meCAR T cells with anti-Fc or meP-647 reagents and showed strong concordance between meCAR expression (anti-Fc) and meP-647 fluorescence. Further, meP-647 did not stain scFv-based CAR T cells, confirming the specificity of meP (**Fig. 2a**).

**Fig. 2:**
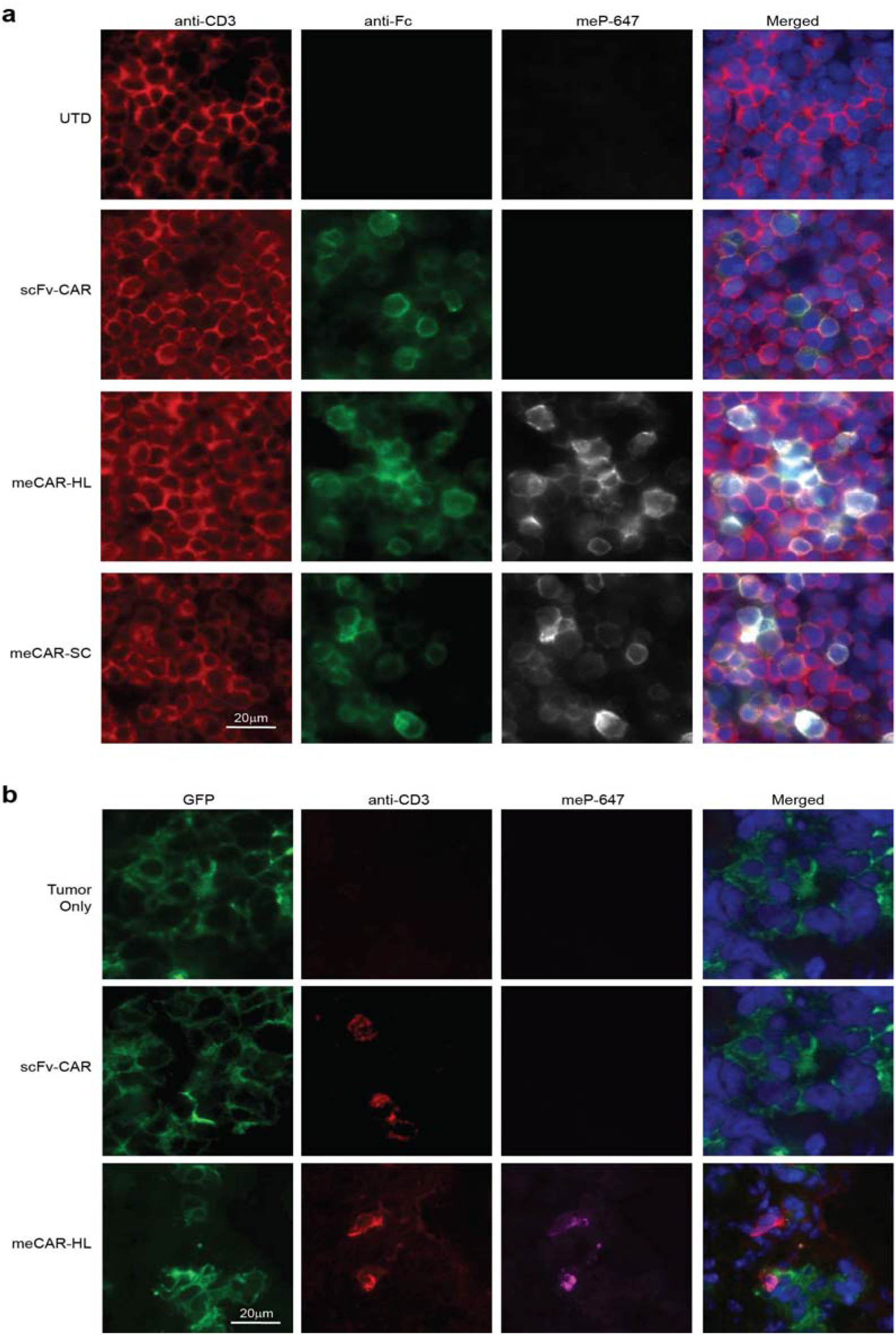
Immunofluorescent Detection of meCAR T Cells using meP-fused fluorophores. **a**, Immunofluorescence staining of untransduced T cells (UTD), HER2 scFv-based HER2-CAR T cells or HER2-meCAR T cells (meCAR-HL or meCAR-SC) stained with anti-CD3, anti-Fc and meP-647. **b**, *In situ* detection of HER2-meCAR T cells in tumor tissue. Subcutaneous GFP+ tumors were injected with either HER2 scFv-based HER2-CAR T cells or HER2-meCAR-HL T cells, and tumors were harvested, fixed and processed for immunofluorescence as in (**a**).

Detection of CAR T cells in patient tissue samples can be technically challenging. To address this, we tested whether meP-647 could be used as a tool to detect meCAR T cells in tumor specimens. We engrafted mice with subcutaneous glioblastoma tumor xenografts, intratumorally injected HER2-meCAR or scFv-CAR T cells, and harvested tumors after 24 hours for analysis. We confirmed the presence of T cells by anti-CD3 immunofluorescence and specifically identified meCAR T cells using meP-647, which co-localized with CD3 staining in tissues treated with meCAR but not scFv-CAR T cells (**Fig. 2b**). Overall, these experiments demonstrated that meP-fused compounds offer a universal strategy for detecting meCAR T cells *in situ* and, thus, could facilitate detection of meCAR T cells in clinical tissue samples.

### Selective expansion of meCAR T cells by meP-fused IL15 cytokine

CAR T cell persistence and expansion following adoptive transfer are critical determinants of therapeutic response^15^. These properties can be enhanced by providing gamma c cytokines, such as IL2 or IL15, either through exogenous delivery or cell engineering^16-18^. IL15 has been shown to support a memory phenotype and enhance CAR T cell function^19, 20^, prompting us to investigate a meP-fused IL15 cytokine for selectively regulating meCAR T cell expansion and persistence. IL15 itself has moderate binding affinity to its beta/gamma chain signaling receptor, and its binding and stability are enhanced when fused to the IL15 receptor alpha sushi domain to generate a chimeric IL15:sushi superagonist (IL15s)^21^. We therefore engineered meP-fused IL15s (meP-IL15s) to evaluate their effects on meCAR T cells (**Fig. 3a**).

**Fig. 3:**
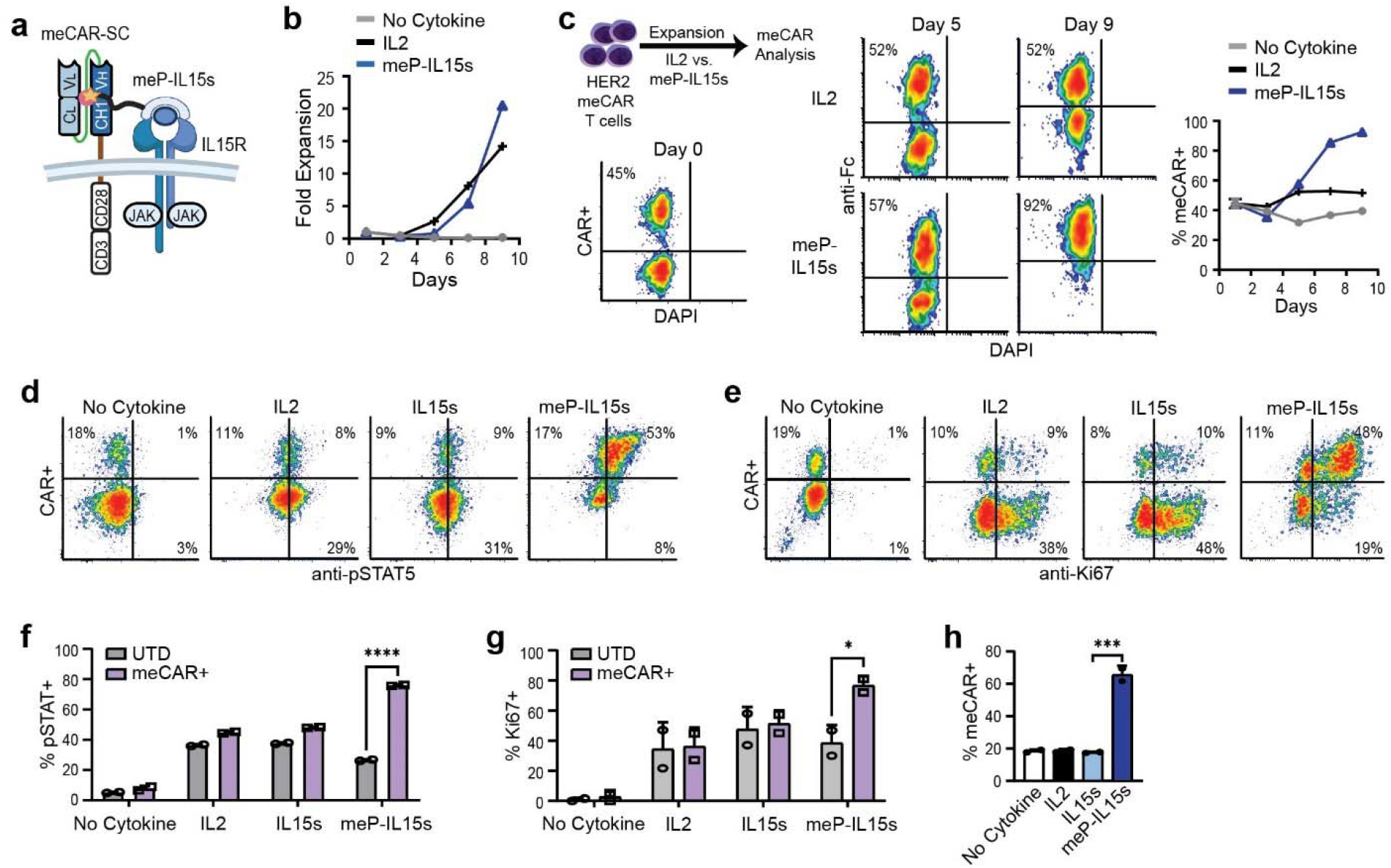
meP-fused IL15s selectively drives meCAR T cell expansion *in vitro*. **a**, Schematic of meP-IL15s interaction with meCAR and IL15-receptor. **b, c**, HER2-meCAR T cells were cultured in IL2 (50IU/mL) or meP-IL15s (20ng/mL) and monitored for expansion (**b**) and the percentage of meCAR+ T cells (**c**) over 9 days. **c**, Representative density plots of DAPI-negative gated cells stained with anti-Fc to detect meCAR+ cells (left), with the percentage of meCAR+ cells in each culture condition over time (right). **d, e**, HER2-meCAR T cells cultured in IL2, IL15s or meP-IL15s for 5 days were stained for CAR expression (anti-Fc) and intracellular pSTAT5 (**d**) or Ki67 (**e**). **f, g**, Mean percentage of viable untransduced (UTD) or meCAR+ T cells that were pSTAT5+ (**f**) and Ki67+ (**g**) after culture with IL2, IL15s, or meP-IL15s (as depicted in the plots of **d, e**). *, P value < 0.05; ****, P value < 0.0001 using a two-way ANOVA test. **h**, Percentages of viable meCAR+ T cells after culture for 5 days with IL2, IL15s or meP-IL15s (i.e., as depicted in the histograms of **d, e**). ***, P value < 0.001 using a one-way ANOVA test.

We cultured HER2-meCAR T cells (45% CAR+) with meP-IL15s or control cytokine IL2 and observed comparable expansion (≥15-fold by day 9; **Fig. 3b**). However, while the percentage of meCAR+ T cells cultured in IL2 remained stable (45-53%), their frequency markedly increased with meP-IL15s, increasing from 57% on day 5 to 92% on day 9 (**Fig. 3c**). This enrichment of meCAR+ T cells by IL15s required the presence of the meP (**Extended Data Fig. 4**). Mechanistically, preferential activation of pSTAT5 and Ki67 were observed in meCAR+T cells cultured with meP-IL15s (**Fig. 3d, e**). In contrast, pSTAT5 and Ki67 levels were comparable between UTD and meCAR T cells under no cytokine, IL2, or IL15s conditions (**Fig. 3f, g**). Notably, meP-IL15s significantly enhanced pSTAT5 and Ki67 expression in meCAR+ T cells and drove their selective expansion within 5 days (from 18% to 66% meCAR+), while other conditions showed no enrichment (**Fig. 3h**). Overall, this data demonstrates that meP-IL15s supports meCAR T cell proliferation and selective expansion of meCAR T cells through *cis* presentation to the IL15 receptor, resulting in pSTAT5 activation.

We next evaluated the utility of IL15s for selectively supporting meCAR T cell persistence and expansion *in vivo*. For these experiments, we administered HER2-meCAR T cells into the peritoneal cavity (i.p.) of NSG mice followed by 10 daily i.p. injections of either IL2 or meP-IL15s (1.0 nmol/mouse) and analyzed the tissue sample on day 9 post treatment (**Fig. 4a**). Treatment with meP-IL15s led to a marked increase in the percentage of meCAR+ T cells in ascites (94%), blood (93%), and spleen (79%) (**Fig. 4b, c**), along with a substantial rise in total meCAR+ T cell numbers across all three tissues (**Fig. 4b, d**). In contrast, control and IL2-treated mice maintained similar levels of meCAR^+^ T cells.

**Fig. 4:**
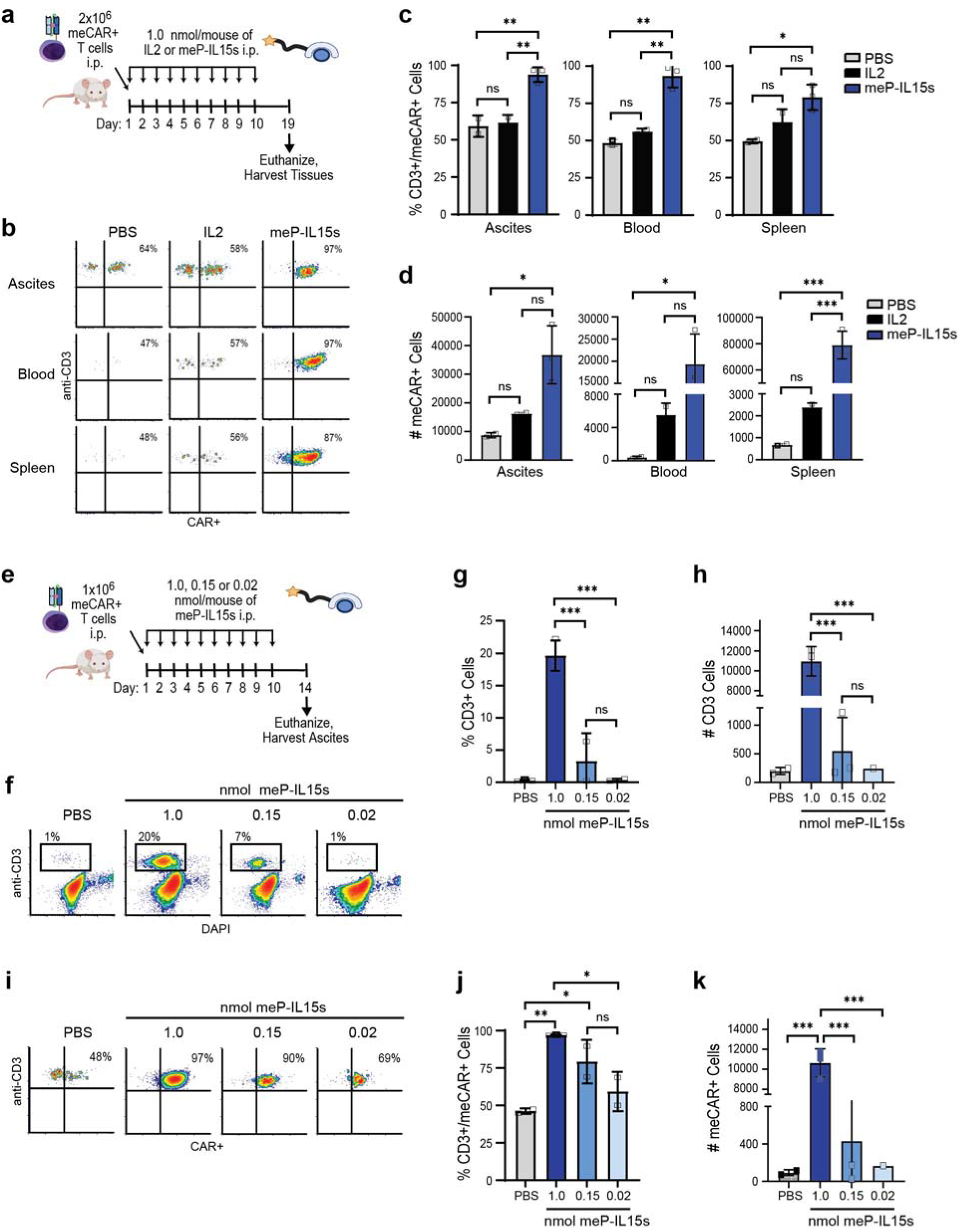
meP-fused IL15s selectively drives meCAR T cell expansion *in vivo*. **a**, Schematic of meP-IL15s and HER2-meCAR T cell treatments *in vivo*. **b**, Tissue samples of ascites, blood and spleen from NSG mice were examined by flow cytometry for the presence of HER2-meCAR+ T cells. Representative dot plots of CD3-gated cells stained with anti-Fc to detect meCAR+ cells. **b-c**, Tissue samples of ascites, blood and spleen were analyzed for the enrichment (**b**) and overall number (**d**) of HER2-meCAR T cells. **e**, Schematic of meP-IL15s dose titration *in vivo* and samples were analyzed on day 14. **f-h**, Ascites samples from NSG mice were examined by flow cytometry (**f**). The enrichment (**g**) and overall CD3+ T cells (**h**) were analyzed at indicated meP-IL15s dosage. **i-j**, The HER2-meCAR T cells were analyzed by meP-647+ cells (**i**) and enrichment (**j**) and overall CD3+ T cells (**k**) were analyzed at indicated meP-IL15s dosage. ***, P value < 0.001; **, P value < 0.01; *, P value < 0.05; ns, non-significant using a one-way ANOVA test. Bars depict mean ± S.D. of the percent cells and numbers of meCAR+ cells in each sample.

To investigate the sensitivity of meCAR T cells to meP-IL15s, we performed a dose titration (0.02-1.0 nmol/mouse) in NSG mouse (**Fig. 4e**). We found the percentage (**Fig. 4f, g**) and the overall CD3+ T cells number (**Fig. 4f, h**) increased at high (1.0 nmol/mouse) meP-IL15 dosage. The lowest dose of meP-IL15s (0.02 nmol/mouse) showed a modest enrichment in meCAR T cells levels from 46% in the PBS condition to 59% meCAR positivity (**Fig. 4i-k**), suggesting that meP-IL15s can selectively enhance persistence of meCAR+ T cells. The intermediate (0.15 nmol/mouse) and high (1.0 nmol/mouse) meP-IL15s dose levels significantly increased meCAR+ T cell levels in a dose-dependent manner (**Fig. 4i-k**). This programmable control is especially important in engineered T cell therapies, where sustained cellular activity correlated with therapeutic efficacy. Together, these data suggest that the meditope platform allows for selective and fine-tune modulation of meCAR T cell persistence and proliferation *in vivo* with post-infusion administration of meP-fused cytokines, such as IL15s.

### Targeting of heterogenous tumors using meCAR T cells and meP-fused antibodies

Tumor antigen heterogeneity remains a major barrier to effective CAR T cell therapy^22^. We hypothesized that the meCAR platform could serve as a universal docking site for meP-fused antibodies, enabling flexible, programmable multiantigen targeting (**Fig. 5a**). To test this, we produced meP-fused anti-CD20 nanobody (meP-αCD20-Nano) and assessed the activity of HER2-meCAR T cells upon co-culture with K562 cells (CD20^-^) or LCL cells (CD20+) with or without meP-αCD20-Nano. While HER2-meCAR T cells did not recognize HER2-negative K562 and LCL cells, the addition of meP-αCD20-Nano induced dose-dependent degranulation as evidenced by CD107a expression on HER2-meCAR T cells co-cultured with CD20+ LCL cells but not CD20-K562 cells (**Fig. 5b**). Moreover, mePαCD20-Nano redirected meHER2-CAR T cell activation (**Fig. 5c**) and killing (**Fig. 5d**) when cultured with LCL cells in a dose-dependent manner. We did not observe activation or killing by scFv-based HER2-CAR T cells with the meP-αCD20, indicating that the effects were meditope-dependent (**Fig. 5c, d**). Similarly, meP-fused αCD20 IgG antibody rituximab (meP-αCD20-IgG) stimulated HER2-meCAR T cells to produce proinflammatory cytokines (GM-CSF, IFNγ, IL-2, and TNFα) when co-cultured with CD20+ Raji tumors but not in the control cultures with UTD T cells or rituximab (**Fig. 5e**). Lastly, the presence of meP-αCD20 antibody did not impact HER2-meCAR T cell killing of HER2+ target cells (**Extended Data Fig. 6b**). These results demonstrate that meP-fused antibodies can effectively and specifically redirect the effector function of meCAR T cells towards alternative tumor antigens, without compromising the recognition of the original CAR target.

**Fig. 5:**
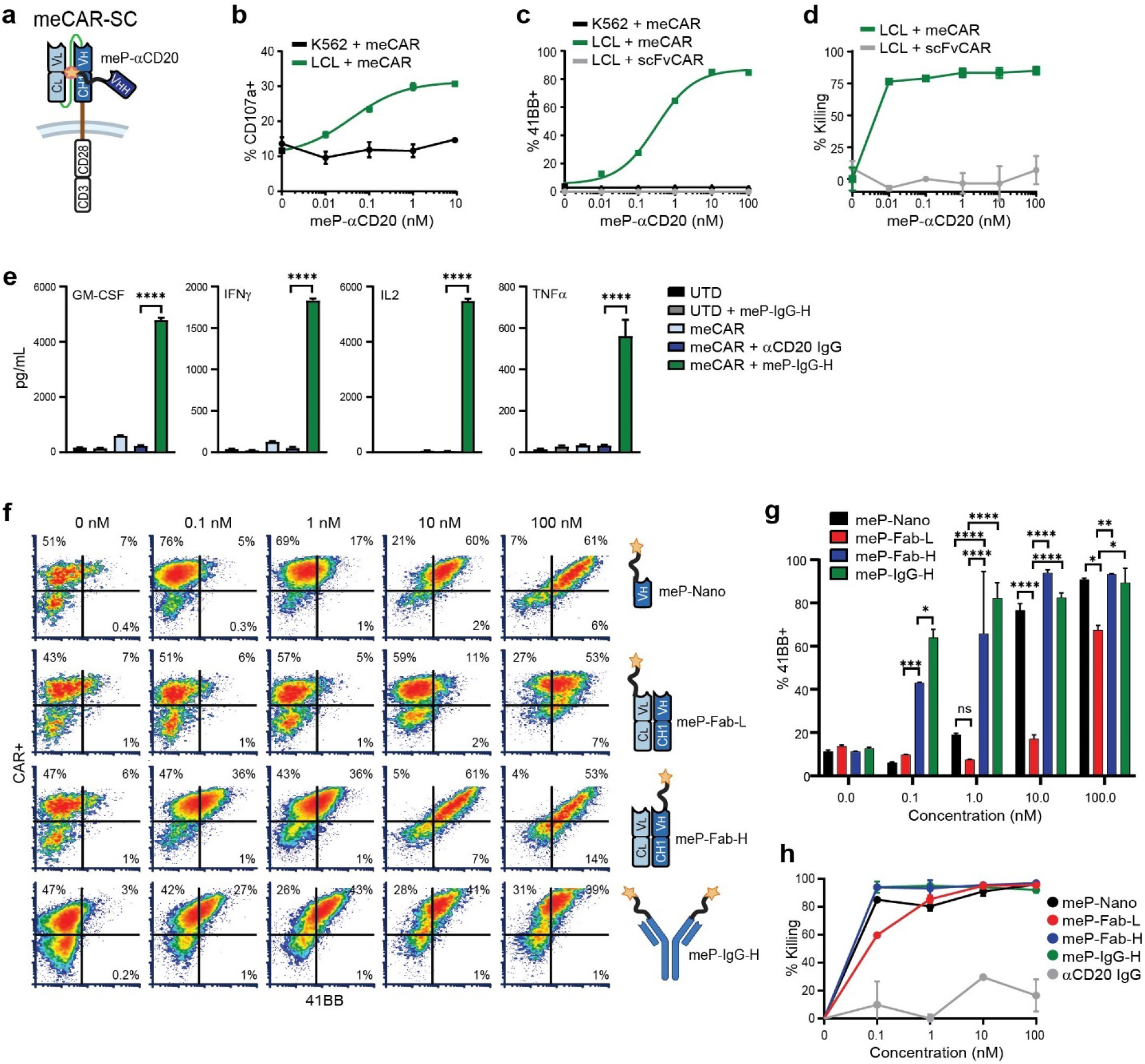
Redirected targeting of HER2 meCAR T cells mediated by meP-αCD20. **a**, Schematic of meP-αCD20 nanobody interaction with meCAR. **b-d**, HER2**-**meCAR T cells were cultured with either CD20-negative K562 cells or CD20+ LCL cells (E:T= 1:1) in the absence or presence of titrated doses of meP-αCD20 nanobody and evaluated by flow cytometry for CD107a expression (**b**), 4-1BB expression (**c**), and tumor cell killing (**d**). Mean ± S.D. of the percentage of the viable meCAR+ gated population (**b, c**) or the viable tumor cell-gated population (**d**) of triplicate wells is depicted. scFv-based HER2-CAR T cells and and CD20-negative K562 cells served as controls. **e**, Cytokine production of HER2-meCAR T cells cultured with Raji tumor cells with or without meP-αCD20 IgG or control αCD20 IgG. UTD T cells were used as the negative control. ****, P value < 0.0001 using a one-way ANOVA test. **f**-**h**, Optimization and functional characterization of meP-αCD20 variants; variant schematics are depicted at right in (**f**). HER2-meCAR T cells were cultured with CD20+ Raji cells with or without meP-αCD20 nanobody, Fab and IgG variants for 48 hours followed by flow cytometric analysis of 4-1BB expression (**f, g**) and tumor cell killing (**h**). **f**, Representative density plots of 4-1BB expression profiles of CD3-gated cells in each culture condition. Mean ± S.D. of the percentage of 4-1BB+ meCAR+ cells (**g**) or tumor killing (**h**) are depicted for each group. ns, nonsignificant; *, P value <0.05; **, P value < 0.01; ***, P value < 0.001; ****, P value < 0.0001 using a two-way ANOVA test.

Since meP-antibodies can bind to either tumor antigens or meCAR T cells, we explored the mechanism of redirection by pre-loading HER2-meCAR T cells or LCL tumor cells with meP-αCD20 followed by washing three times before co-culture with the unlabeled counterpart cells (**Extended Data Fig. 5a**). We observed a more significant increase of CD107a expression on meCAR T cells in the tumor-bound (77%) compared to the T cell-bound condition (16%) (**Extended Data Fig. 5b**), suggesting that redirected targeting of meCAR T cells by meP-antibodies occurs primarily through pre-binding of meP-antibodies to tumor cells, which are then recognized by meCAR T cells.

We next compared antibody formats for meP-αCD20, testing nanobody, Fab, and IgG variants with meP fused at the N-terminus of either the heavy or light chain (**Fig. 5f**). All formats redirected HER2-meCAR T cell 4-1BB activation (**Fig. 5f, g**) and killing (**Fig. 5h**) against CD20+ Raji cells, although the meP-Fab-L variant was less effective at low concentrations (**Fig. 5f-h**). Redirected meCAR T cell activation and killing were meP-dependent, as they were not observed using the control αCD20 antibody lacking the meP (rituximab) (**Fig. 5h, Extended Data Fig. 6a**). These findings demonstrate the versatility of the meCAR platform in accommodating diverse antibody formats, including nanobody, Fab and IgG, for tumor antigen redirection. The results also suggest that N-terminal heavy chain fusion is optimal for redirection and will be the default format unless otherwise noted.

To explore the utility of meP-antibodies to redirect meCAR T cell specificity *in vivo*, NSG mice were engrafted i.p. with HER2-negative CD20-engineered MDA468 (MDA468-CD20) tumors and treated with HER2-meCAR T cells and meP-αCD20 IgG every two days for 10 doses (**Fig. 6a**). Mice treated with the combination of meP-αCD20 IgG and HER2-meCAR T cells exhibited greater tumor reduction (**Fig. 6b**) and extended survival (**Fig. 6c**) when compared to mice treated with meCAR T cells alone or combined with αCD20 antibody lacking meP (rituximab).

**Fig. 6:**
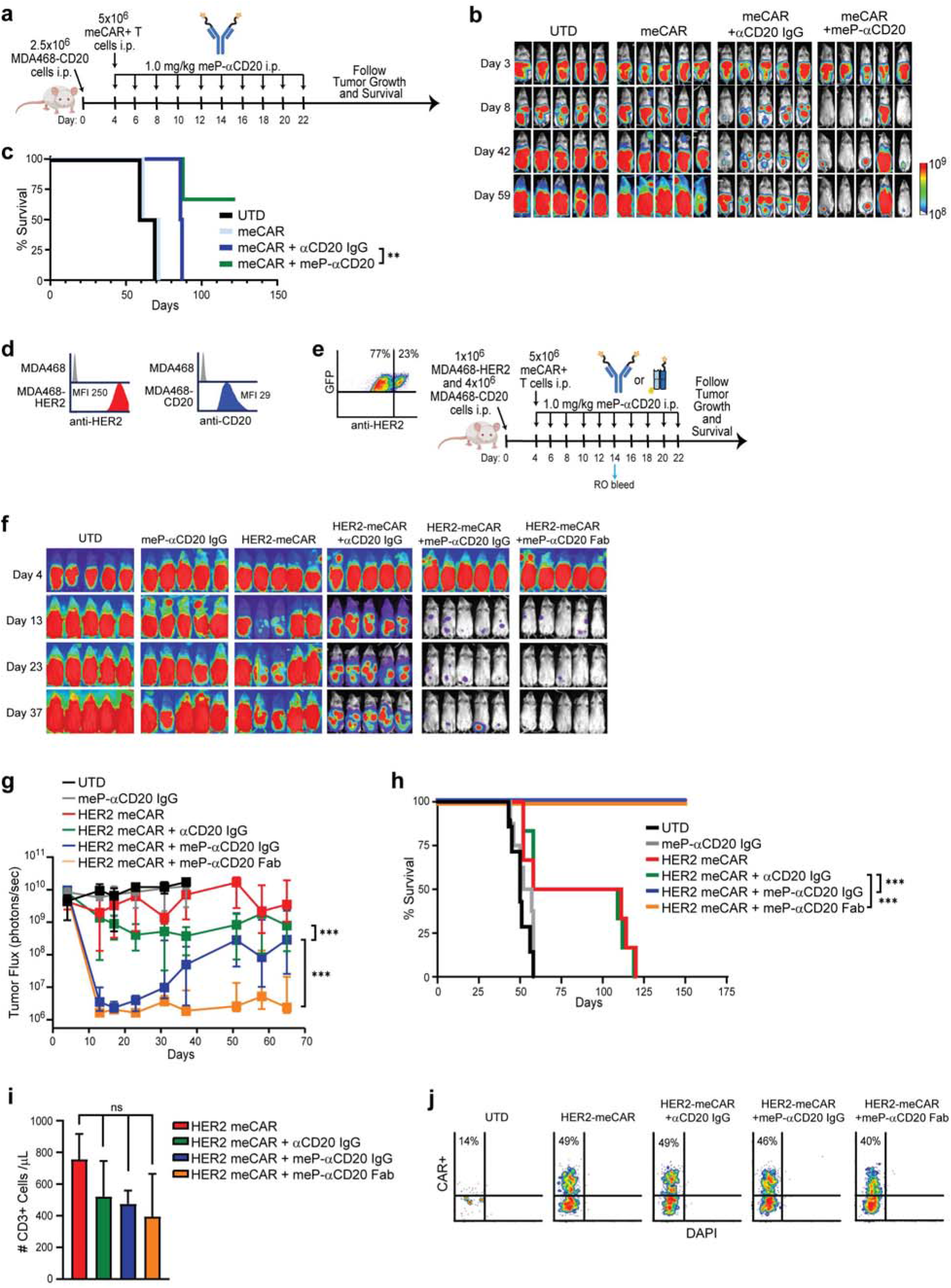
Dual targeting of HER2-meCAR T cells mediated by meP-αCD20 in a heterogenous tumor model. **a-b**, Schematic of redirected targeting of HER2-meCAR T cells *in vivo* (**a**). NSG mice bearing ffLuc+ MDA468 tumors engineered to express CD20 (MDA468-CD20) were treated with HER2-meCAR CAR T cells, with or without meP-αCD20 (meP-IgG-H variant), followed by biophotonic imaging for tumor size over time. Mice treated with UTD T cells were used as the negative control (**b**). **c**, Kaplan Meier survival analysis of each group. **, P value < 0.01 using a Mantel-Cox test. **d-e** Schematic of dual targeting of HER2-meCAR T cells *in vivo*. ffLuc+ MDA468 tumor cells engineered to express CD20 or HER2 cells (**d**) were grafted to mimic tumoral heterogeneity (**e**) followed by HER2-meCAR T cells with or without αCD20 reagent (control αCD20 IgG, meP-αCD20 IgG or meP-αCD20 Fab). **f**, Tumor growth was monitored by biophotonic imaging. **g**, Mean ± S.D. tumor flux value for each group is depicted. ***, P value < 0.001 using a two-way ANOVA test. **h**, Kaplan Meier survival analysis of each group. ***, P value < 0.001 using a Mantel-Cox test. **i-j**, Blood samples collected on day 14 were evaluated for total T cells (**i**) and HER2-meCAR T cells by flow cytometry (**j**). ns, nonsignificant, using a one-way ANOVA test.

The Fc region of IgG supports serum stabilization and extended *in vivo* half-life, which is critical for therapeutic efficacy *in vivo*. However, the small size of Fab reagents offers advantages, including better tissue penetration and reduced steric hindrance. As our pilot studies using prototype meP-αCD20 Fab showed a lack of therapeutic efficacy (**Extended Data Fig. 6c**), we used mouse serum albumin peptide (MSAP) to stabilize and increase *in vivo* half-life. We determined that fusing MSAP to the C-terminus of either the light chain (LCC) or heavy chain (HCC) of the Fab did not interfere with the meP-αCD20 Fab redirected killing or IFNγ production by HER2-meCAR T cells (**Extended Data Fig. 6d, e**). Mice engrafted with MDA468-CD20 and treated with MSAP-fused meP-αCD20 Fab and HER2-meCAR T cells exhibited tumor reduction (**Extended Data Fig. 6f, g**). In this model, the LCC variant demonstrated better *in vivo* efficacy than the HCC variant (**Extended Data Fig. 6f, g**), and we used the meP-αCD20 Fab LCC format in subsequent experiments. Together, these studies highlight the feasibility of the meCAR platform to redirect T cell specificity *in vivo* using meP-fused antibodies and underscore the importance of optimizing platform features, such as meP-fusion position and antibody half-life, for maximal therapeutic efficacy.

We next evaluated the meCAR platform as a strategy to overcome tumor heterogeneity by developing a mixed-antigen solid tumor model expressing both CD20 and HER2 (**Fig. 6d**). For these studies, we engineered GFP/ffLuc+ MDA468 cells to express either HER2 (MDA468-HER2) or CD20 (MDA468-CD20) and i.p. engrafted the tumor cells at a 1:4 ratio (MDA468-HER2:MDA468-CD20) in NSG mice, which yielded tumors with mixed antigen expression (**Fig. 6e**). Upon treatment with HER2-meCAR T cells alone or with control αCD20, we observed partial responses (**Fig. 6f-h**), with significant reduction of tumor burden and improved survival only seen upon treatment with HER2-meCAR T cells combined with meP-αCD20 IgG or Fab (**Fig. 6f-h**). Examination of circulating T cells in the blood at day 14 revealed comparable levels of HER2-meCAR T cells in all groups (**Fig. 6i, h**). Together, these results illustrate the dual-targeting flexibility of meCAR T cells combined with meP-fused Fab, both in terms of antigens and tumor models.

## Discussion

CAR T cell therapies have demonstrated transformative clinical activity, particularly in hematological malignancies^2, 23, 24^. However, several challenges continue to limit their broader applicability, including fixed antigen specificity, limited ability to regulate cell activity post adoptive transfer, and insufficient persistence and functionality in the immunosuppressive TME^25, 26^. These limitations are especially pronounced in solid tumors but also impact the potency and safety across cancer types. To overcome these barriers, we developed a meditope-based, plug-and-play CAR T cell platform that enables precise and modular functional enhancement without altering the underlying CAR specificity. Our proof-of-concept studies demonstrate that meditope-binding can be readily incorporated into Fab-based CAR constructs of diverse specificities and leveraged to introduce programmable features, such as selective expansion of CAR-engineered T cells both *in vitro* and *in vivo* using a meP-fused cytokine and tunable dual-antigen targeting via meP-fused antibodies. The meCAR/meP platform is not limited to CAR antigen specificity, tumor type, or meP conjugate and, therefore, may be broadly and rapidly applied across the CAR T cell field.

The extent of CAR T cell expansion and persistence following infusion strongly impacts therapeutic efficacy. One strategy to enhance these properties involved co-expression of gamma chain cytokines, such as IL-2 and IL-15, resulting in so-called ‘armored’ CARs^27, 28^. A recent study reported superior tumor control by GPC3-targeted CAR T cells that co-expressed IL-15 than CAR T cells without IL-15 in patients with various solid tumors, highlighting the utility of armored CARs^29^. However, these approaches hardwire cytokine production into the CAR design, limiting the ability to modulate activity, and raising concerns about altered phenotypes during manufacturing or increased toxicity post-infusion. Here we demonstrate that meP-IL15s, a soluble cytokine fused to the meditope peptide, selectively enhances expansion and persistence of meCAR T cells in a dose-dependent manner, offering a modular and titratable system to support CAR T cell function. This approach bypasses the need for additional transgenes, simplifying manufacturing and reducing the risk of uncontrolled cytokine-driven expansion. Mechanistically, meP-IL15s selectively activates STAT5 signaling and proliferation in meCAR+ T cells through cis engagement, confirming the specificity of the interaction. Importantly, this external control system enables dynamic regulation of CAR T cell activity post-infusion, a feature that could be tailored to patient needs or disease context. The meP-based delivery system may be broadly adapted to deliver a range of cytokines or modulators, extending its potential as a flexible tool to optimize CAR T cell performance.

Tumor heterogeneity remains a major challenge in CAR T cell therapy, particularly in the context of solid tumors. Although single antigen-targeted CARs have shown remarkable clinical success in specific B-cell malignancies^1-3, 5^, multi-antigen targeting approaches are likely required to address tumor heterogeneity. Several groups have developed bispecific CARs with dual-antigen specificity, which have shown promising preclinical and clinical activity across several tumor types^30-38^. However, bispecific CARs are inherently limited by their fixed antigen specificity, making it challenging to adapt to progressive antigen escape and immunoediting. In contrast, CAR platforms that incorporate antibody-based switchable elements provide flexibility in antigen targeting, enabling modification of the targeting strategy during the course of treatment based on the dynamics of tumor antigen expression^39-41^. Here we show that meP-fused antibody variants, including IgG, Fab, and nanobody, can effectively redirect meCAR T cells against alternative tumor antigens *in vitro* and *in vivo*, while preserving engagement of the original target. The meCAR platform integrates the strengths of both hard-wired and flexible designs, with the meCAR T cells recognizing the predominantly expressed tumor antigen and the meP-antibodies mediating the targeting of additional tumor antigens that can be adapted based on tumor type or tumor evolution. This plug-and-play system provides a versatile solution to tumor heterogeneity, allowing multi-antigen targeting without the need for re-engineering or re-manufacturing of CAR T cells.

In summary, our meCAR platform leverages the meditope technology to confer versatile functionality to CAR T cells, offering new opportunities for imaging, manufacturing, in vivo expansion, and multi-antigen targeting. While the studies presented here serve as proof-of-concept, our findings support the broader applicability of this approach to CARs targeting diverse antigens and to a wide array meP-fused biologics. This platform provides a flexible framework to address several key limitations of current CAR T cell therapies, particularly in the context of solid tumors. Looking forward, we anticipate that the meCAR platform can extended to incorporate additional functionalities, including modifiers of the TME to overcome immunosuppression, safety switches to enable enhanced control, and labeled conjugates to facilitate dynamic imaging of CAR T cells.

### Online Methods

#### Cell lines

Human breast cancer cell lines MCF7 (ATCC HTB-22), SKBR-3 (ATCC HTB-30) and HT-1080 (ATCC CCL-121) were maintained in DMEM (Gibco #11960051) supplemented with 10% heat-inactivated FBS (Hyclone) and 2 mmol/L L-glutamine (Irvine Scientific). Lymphoblastoid cell lines (LCL) and Raji (ATCC CCL-86) were maintained in RPMI (Gibco #22400089) supplemented with 10% heat-inactivated FBS (Hyclone). To generate tumor cell lines expressing enhanced green fluorescent protein (GFP) and firefly luciferase (ffLuc), MCF7, Raji, OVCAR3 and MDA468 cells were transduced with a lentiviral vector encoding GFP-ffLuc. After expansion in complete medium for 14 days, GFP+ cells were sorted by FACS for >98% purity and expanded for *in vivo* experiments.

#### Meditope-enabled CAR (meCAR) constructs and lentivirus production

The clinical grade scFv-based HER2-meCAR was generated as previously described and the Fab region used in the meHER2 CAR construct was obtained from meditope-enabled trastuzumab^8, 11, 42^. Briefly, the expression of VL/CL and VH/CH were mediated by a T2A ribosomal skip mechanism to generate HER2 meCAR-LH or meCAR-HL, depending on the expression order of VL/CL and VH/CH (**Fig. 1a**). Additionally, we utilized the L60 linker to stabilize the pairing of VL/CL and VH/CH for creating the single-chained format of the HER2 meCAR-SC construct. The CD19- and HER3/EGFR meCARs were created by CDR grafting from meditope-enabled FMC63 and MEHD7945A monoclonal antibodies, respectively, to a 4D5 framework (**Extended Data Fig. 1a**)^1, 14^. The extracellular domain included 129 amino acids of hinge, linker and CH2-deleted IgG4 Fc (CH3) and the intracellular domains were composed of CD28 co-stimulatory and cytolytic CD3ζ signaling domains^42, 43^. 293T cells were transfected with meCAR constructs and lentiviral packaging plasmids using polyethylenimine (PEI) methods. Supernatants were collected and followed by high-speed centrifugation to enrich lentivirus. Viral titers were determined using Jurkat cells based on Fc expression.

#### CAR T cell production

Naïve/memory T cells (Tn/mem) were isolated from healthy blood donors under protocols approved by City of Hope (COH) Institutional Review Board^35, 44^. Briefly, peripheral blood mononuclear cells (PBMCs) were isolated by density gradient centrifugation over Ficoll-Paque (GE Healthcare) and underwent a series of selection by CliniMACS/AutoMACS to remove CD14+ and CD25+ cells, followed by CD62L positive selection for Tn/mem cells. To generate CAR T cell products, T cells were stimulated with Dynabeads Human T-Expander CD3/CD28 (Invitrogen) at a 1:3 ratio (T-cell:bead) overnight in X-VIVO-15 (Lonza) supplemented with 10% FBS, 2 mmol/L L-glutamine, 50 U/mL IL-2 and 0.5 ng/mL IL-15. Stimulated T cells were then transduced with lentivirus (MOI of 1.0) encoding the meCAR. Mock and CAR transduced T cells were cultured with indicated cytokines three times a week for 16-18 days before subsequent analyses.

#### Animal studies

All mouse experiments were approved by the COH Institutional Animal Care and Use Committee (IACUC protocol #18059). For MCF7 and OVCAR3 models, the NSG mice were intraperitoneally (i.p.) injected with 5×10^6^ of ffLuc+ tumor cell for 7 days, followed by randomized grouping for i.p. treatments with 2×10^6^ of UTD or CAR T cells on day 8. For redirected and dual-targeting models, the NSG mice were intraperitoneally (i.p.) injected with ffLuc+ MDA468-CD20 or mixture MDA468-CD20/MDA468-HER2 tumor cells, as indicated in the experimental schema for 4 days, followed by randomized grouping for i.p. treatments with 5×10^6^ of UTD or meCAR T cells. Mice were then treated with meP-αCD20 antibodies at a dose of 1.0mg/Kg every 2 days for a total of 10 administration. Tumor burden was imaged at the indicated time point with SPECTRAL LagoX (Spectral Instruments Imaging) and analyzed using Aura software (v2.3.1, Spectral Instruments Imaging). Survival curves were generated by GraphPad Prism Software. Tissue harvesting for evaluation of tumor antigen expression in end stage mice occurred after euthanasia.

#### Generation of meP-fused compounds

IL15superagonist: Fc-IL15superagonist and Fc-meP-IL15superagonist fusion were transiently expressed in a Chinese hamster ovary (CHO) cell line using the ExpiCHO™ system (Thermo Fisher Scientific) according to the manufacturer’s manual. The culture supernatant was collected at day 7 and purified through affinity chromatography (ProG resin, Genscript) followed by size exclusion chromatography (Cytiva Lifesciences). To generate IL15superagonist (IL15s) and meP-IL15superagonist (meP-IL15s), Fc was cleaved through the TEV cleavage site between Fc and IL15superagonist with and without meP and removed through reverse affinity chromatography. Biophysical validation of those molecules was performed via Surface Plasmon Resonance (SPR, Biacore T200, Cytiva Lifesciences).

Rituximab and its meP-fused Fabs and IgGs through N-terminus of heavy chain or light chain were produced in the ExpiCHO-S™ cells (Thermo Fisher Scientific). After transfecting the expression vectors for rituximab or variants using ExpiFectamine™ (Thermo Fisher Scientific), cell cultures were incubated in a 37°C shaking incubator with >80% humidity and 8% CO_2_ for 1 day and then moved to a 32°C shaking incubator with 5% CO_2_ for the next 6 days. Purification was done with the appropriate affinity and size exclusion chromatography. Binding affinity for antigen and meditope-enabled antibody was measured on Biacore T200 (Cytiva Lifesciences). Kinetic characterization was determined by Biacore evaluation software.

#### Flow cytometry

Expression of meCARs on T cells was analyzed with meP-647 (City of Hope) or antibodies against human Fc (Jackson ImmunoResearch cat# 109-096-008), and clinical scFv CAR expression was determined by staining truncated CD19 (BD Biosciences). T cell activation was determined using antibodies against CD107a, 4-1BB and IFNγ (BD Biosciences). Antibodies for pSTAT5 (BD Biosciences clone 47) and Ki-67 (Biolegend clone 16A8) were used to analyze preferential meCAR expansion by meP-IL15s. All samples were analyzed via a MACSQuant Analyzer (Miltenyi Biotec Inc.) and processed via FCS Express 7.

#### In vitro T cell functional assays

Tumor cells were cultured in the medium described above, and meCAR T cells were washed/resuspended in the same medium followed by co-culturing with tumor cells at indicated Effector:Tumor (E:T) ratio. For degranulation and intracellular IFNγ assays, meCAR and tumor cells were incubated at an E:T of 1:1 for 5h with the addition of anti-CD107a (BD Biosciences) and GolgiStop protein transport inhibitor (BD Biosciences). After the coculture, cells were harvested, fixed, permeabilized, and analyzed by flow cytometry as described above. To test cytolytic activity, tumor and CAR+ T cells were co-cultured at indicated E:T ratios for 48h and the killing was quantified by flow cytometry analyzing viable tumor cells (GFP+/DAPI-).

#### In situ Immunofluorescence staining

Primary brain tumor (PBT) PBT103-2 were obtained from GBM patient resections at COH under protocols approved by the COH IRB^45^. 1.0×10^6^ of tumor cells were subcutaneously engrafted in NSG mice until tumor volume reaches 100 mm^3^ followed by intratumorally treatments of 2×10^6^ of scFv or meCAR T cells. GBM tumors were frozen sectioned and fixed in 4% paraformadehyde for 15 minutes. After fixation, samples were blocked in PBS containing 0.25% BSA and 0.3% Trixton X-100 for 1 hour. Primary antibodies of rabbit anti-human Her2 (Agilent A048529-1) and mouse anti-human CD3 (Agilent M725429-2) antibodies were used for 1 hour followed by anti-rabbit Alexa 488 (ThermoFisher A-21206), anti-mouse Alexa 555 (ThermoFisher A-31570) and meP-647 staining for 30 minutes. Finally, samples were counter stained by DAPI.

#### Statistical analysis

Data were presented as mean ± SD and statistical comparisons were analyzed using the one-way or two-way ANOVA for independent or grouped samples to calculate p-value, unless otherwise stated. *p < 0.05, NS, not significant. Comparison of Kaplan-Meier survival data was performed using Prism v9.0 (GraphPad Software).

#### Cytokine profiling

HER2-meCAR T cells were cultured with Raji tumor cells (E:T= 1:1) in the absence or presence of either meP-αCD20 IgG or control αCD20 IgG (100nM) for 48 hours. Harvested supernatants were analyzed for GM-CSF, IFNγ, IL2 and TNFα by cytometric bead array using the Human Cytokine 10-Plex Panel (ThermoFisher) according to the manufacturer’s instructions.

## Acknowledgements

We thank Alfonso Brito and Yuelong Ma for their technical assistance with this study. We also thank the current and former members of the Brown, Forman and Williams group for helpful discussions. This study was funded by grants from NCI R21-CA193055 (J.C.W., C.E.B.) the Marcus Foundation (C.E.B., J.C.W.) and City of Hope IDDV program. We also thank Core Facilities at City of Hope, including Small Animal Imaging and Drug Discovery and Structural Biology Share Resources supported by NCI grant P30CA033572.

## Author Contributions

CK, ZT, SJF, JCW and CEB conceptualized the research.CK, ZT, YK, LS, ZW, BA, DA, RS and WC design and collect data. CK, ZT, and YK analyze results. MP, JK, KL, BP and JCW expressed and design meP-fused compounds. CK, JRO, MCC, DR, SJF, JCW and CEB help manuscript preparation. SJF, JCW and CEB supervised the research.

## Competing Interests

C.E.B. and S.J.F. report personal fees, patent royalties and research support from Chimeric Therapeutics during the conduct of the study, as well as patent royalties from Mustang Bio outside the submitted work. J.C.W. reports personal fees, patent royalties and research support from Xilio Tx and Meditope Biosciences. All other authors declare no competing interests

