## Supplemental figures for "Meditope-Enabled Chimeric Antigen Receptors Facilitate Plug-and-Play Control of T Cells"

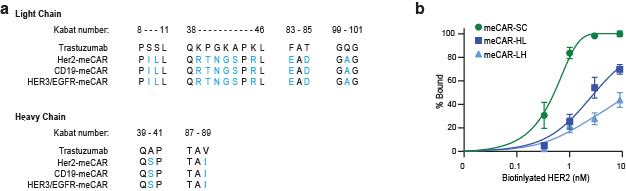
**Extended Data Fig. 1|** **Amino acid substitution for meCAR design and affinity characterization of meCAR varianats. a.** Relevant light chain (top) and heavy chain (bottom) amino acid sequences in accordance with the kabat number of trastuzumab as compared to the HER2-, CD19- and HER3/EGFR-meCARs. The amino acids that interact with meP are highlighted in blue. **b.** Affinity titration of HER2-meCAR variants. Jurkat cells were transduced to stably express HER2-meCAR-SC, -HL, or -LH variants and incubated with biotinylated HER2 antigen followed by SA-PE staining. The percentage of HER2 antigen binding was based on meCAR+ cells as detected with meP-647.

**Extended Data Fig. 2| *in vivo* efficacy of HER2 meCAR T cells in an OVCAR-3 xenograft model. a.** Characterization of HER2 antigen expression in moderately expressing MCF7 and highly expressing SKBR3 tumor cells. Grey histogram indicated isotype control; MFI, mean fluorescence intensity. **b.** ffLuc+ OVCAR-3 cells were intraperitoneally injected in NSG mice (5x10^6^ cells/mouse). On day 8 post-implantation, mice were i.p. treated with either UTD T cells, HER2 meCAR T cells or HER2 scFvCAR T cells (2x10^6^ cells/mouse) followed by biophotonic imaging for tumor size over time **c.** Tumor burden and survival analysis tumor-bearing mice. Bars depict mean ± S.D. flux values for each group. *, P value < 0.05; ns, nonsignificant using a two-way ANOVA test. **d.** Kaplan Meier survival analysis of each group. Dashed line indicates day of T cell treatment. ***, P value < 0.001; ns, non-significant using a Mantel-Cox test.

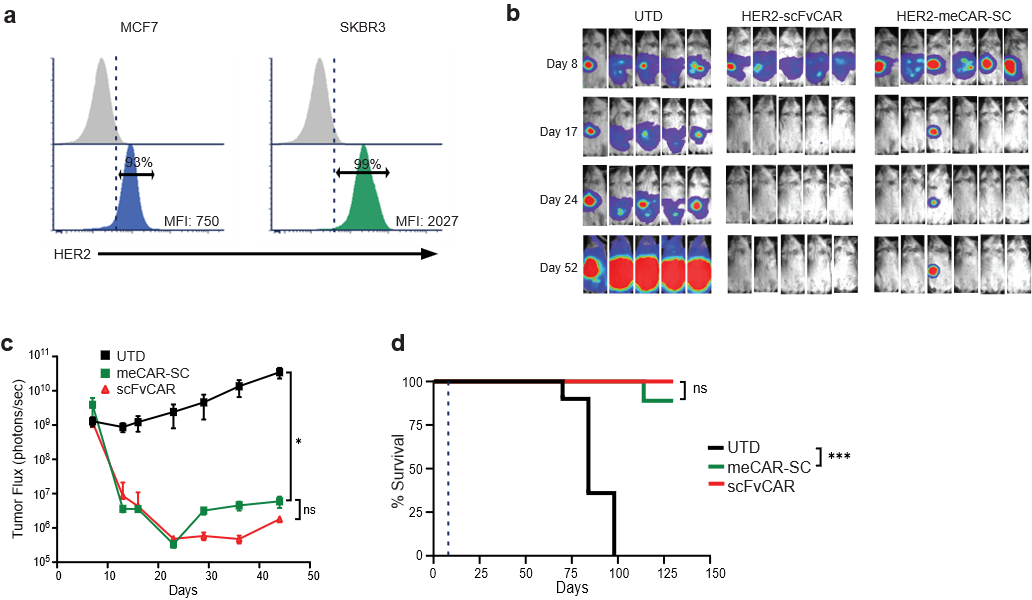

**Extended Data Fig. 3| Design and characterization of meditope-enabled CARs targeting additional antigens. a**, Design, expression and meP binding of humanized CD19-meCAR T cells. CD19-scFvCAR vs. CD19-meCAR expression was characterized by staining with anti-Fc (left), and meP-647 (right) followed by flow cytometry. Untransduced (UTD) T cells were used as controls. **b-d,** CD19-meCAR and scFvCAR T cells were cultured with Raji tumor cells and analyzed by flow cytometry for viable tumor cell number (**b**), percentages of CAR+ T cell expansion (**c**), or percentages of Raji cell cytolysis at a range of E:T ratios (**d**). Bars depict mean ± S.D. from duplicate samples. **e**, HER3/EGFR-meCAR expression was characterized by staining with anti-Fc (left), and meP-647 (right) followed by flow cytometry. UTD T cells were used as controls. **f**, Characterization of HER3 and EGFR expression on MCF7 and SKBR3 tumor cells. Grey histograms represent isotype control. **g**, Kinetics of tumor cell killing performed by xCELLigence assay upon co-culture of the HER3/EGFR-meCAR T cells with either MCF7 (top) or SKBR3 (bottom) tumor cells in the absence (blue line) or presence (red line) of meP-647 at 100nM.

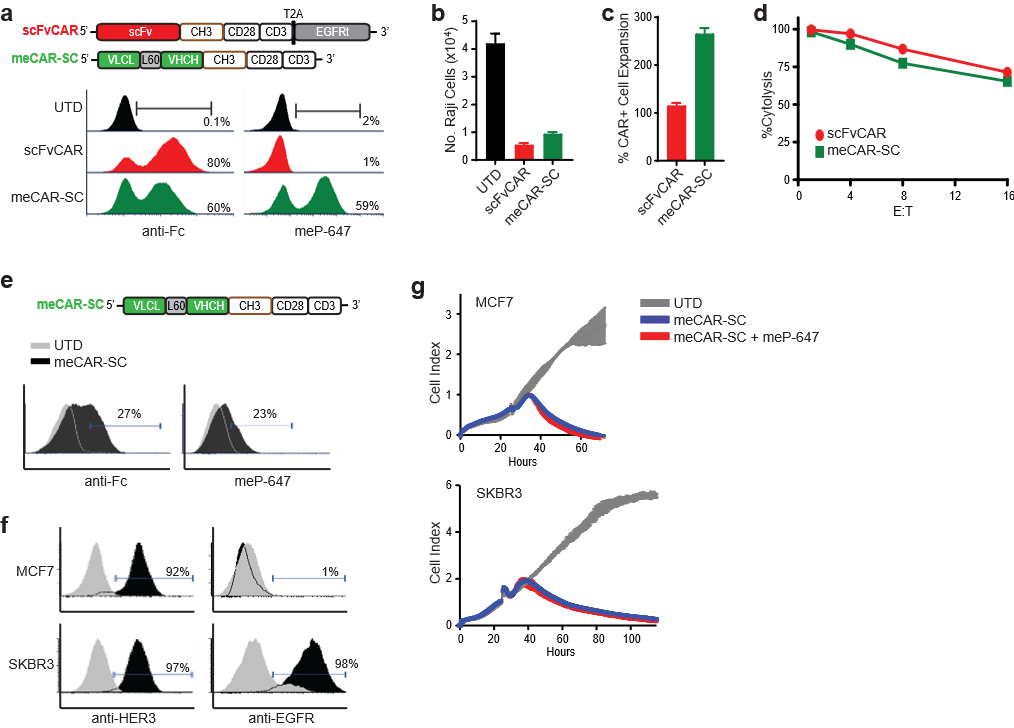

**Extended Data Fig.4| Manufacture of HER2 meCAR T cells with meP-IL15s. a,** Schematic of HER2-meCAR T cell production. Briefly, Tn/mem cells were stimulated with CD3/CD28 beads followed by lentiviral transduction on day 1. Medium was replenished every 2 days with cytokine supplementation described in methods. On day 7, HER2-meCAR T cells underwent bead removal, and were split into cultures with either IL-2, IL15s or meP-IL15s and expanded for an additional 9 days. **b,** HER2-meCAR T cellswere analyzed for meCAR+ T cell enrichment on day 10, 12 and 16 using the anti-Fc antibody.

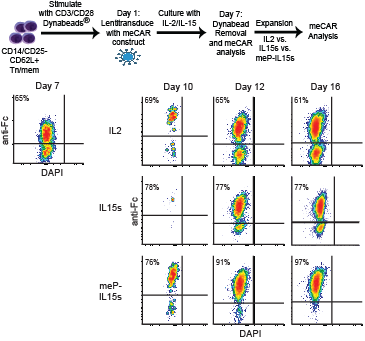

**Extended Data Fig. 5| Binding of meP-αCD20 on tumor triggers dual-targeting of HER2 meCAR T cells. a,** Schematic of “tumor bound” and “T cell bound” conditions for dual-targeting assay. Briefly, LCL (CD20+) cells (top) or HER2-meCAR T cells (bottom) were incubated with meP-αCD20, and washed 3 times to remove excessive meP-αCD20 prior to initiation of co-culturesat anE:T of 1:1. **b,** Quantitation of CD107a induction after 5 hours of co-culture. Reperesentative histograms are depicted at left. Percentages of CD107a-expressing cells in the meCAR+ gated population (based on anti-Fc staining) as compared to CD107a expression on UTD cells are shown at right. Bars depict mean ± S.D. from duplicate wells.

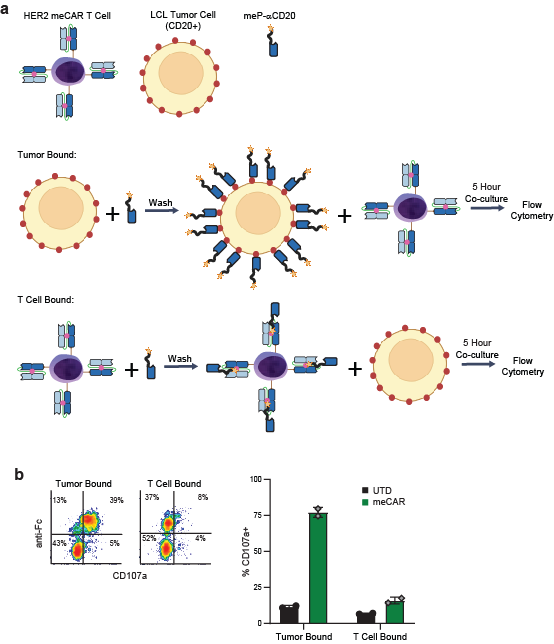

**Extended Data Fig. 6| Optimized meP-αCD20 Fab shows improved antitumor activity. a,** Raji cells were co-cultured with HER2-meCAR T cells in the presence (100nM) of control αCD20 IgG for 48 hours followed by flow cytometry analysis of 4-1BB activation. **b,** MCF7 cells were co-cultured with UTD or HER2-meCAR T cells in the presence (100nM) or absence of meP-αCD20 IgG for 48 hours followed by killing analysis. *, P value <0.05; ns, nonsignificant using one-way ANOVA test. **c,** Raji tumor cells were intraperitoneally (i.p.) injected into NSG mice (5x10^5^ cells/mouse). On day 4 post-implantation, mice were i.p. treated with HER2-meCAR T alone or in combination with meP-αCD20 Fab followed by biophotonic imaging for tumor size over time. **d-e,**  The mouse serum albumin peptides (MSAP) were fused to C-terminus of light chain (LCC) or heavy chain (HCC) of meP-αCD20 fab. HER2**-**meCAR T cells were co-cultured with Raji cells at an E:T of 1:1 for 48 hours with titrated doses (0 to 100nM) of either meP-αCD20 Fab, meP-αCD20 Fab with MSAP fused to the C-terminus of either the light chain (LCC) or heavy chain (HCC), or negative control αCD20 IgG, and evaluated by flow cytometry for killing (**d**) and IFNγ production (**e**). **f-g,**  ffLuc+ MDA468-CD20 tumor cells were i.p. injected into NSG mice (2.5x10^6^cells/mouse). On day 4 post-implantation, mice were treated with an i.p. injection of either UTD or HER2-meCAR CAR T cells (5x10^6^ cells/mouse) with or without meP-αCD20 (LCC or HCC variants, 1.0mg/kg) every 2 days for 10 doses, followed by biophotonic imaging for tumor size over time.

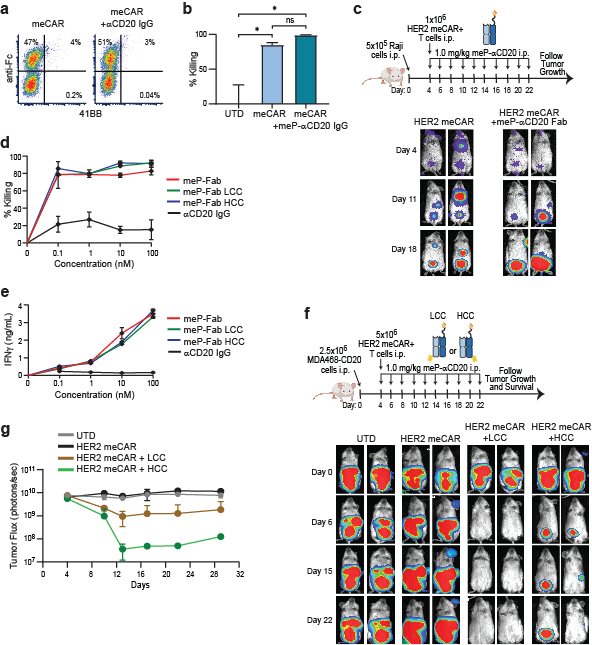
